# Navigating the Cold: Integrative Transcriptome Sequencing Approach Reveals Ionoregulatory and Whole-Body Responses to Cold Acclimation in *Drosophila ananassae*

**DOI:** 10.1101/2024.07.18.604044

**Authors:** Vera Miyase Yılmaz, Zhihui Bao, Sonja Grath

## Abstract

Understanding how species adapt to changing environments is a major goal in evolutionary biology and can elucidate the impact of climate change. Climate imposes inevitable effects on the geographical distribution of insects as their body temperature primarily depends on the environment. The vinegar fly *Drosophila ananassae* expanded from its tropical ancestral range to more temperate regions, which requires adaptation to colder temperatures. Transcriptome and genome-wide association studies focusing on the ancestral-range population identified the targets of selection related to ionoregulatory tissues. However, how cosmopolitan *D. ananassae* adapted to colder environments, where low temperatures last longer, is still unknown. Here, we present a study on the effect of long-term cold exposure on *D. ananassae*, examining the gene expression variation in the whole body and the ionoregulatory tissues, namely the hindgut and the Malpighian tubule. To elucidate molecular mechanisms of cold adaptation during species expansion, we included cold-tolerant and cold-sensitive strains from the ancestral species range and cold-tolerant strains from the derived species range. We show that cold acclimation improves cold tolerance and results in differential expression of more than half of the transcriptome in the ionoregulatory tissues and the whole body. Notably, we provide complementary insight into molecular processes at four levels: strains, populations, phenotypes, and tissues. By determining the biochemical pathways of phenotypic plasticity underlying cold tolerance, our results enhance our understanding of how environmental changes affect thermal adaptation in natural populations.

**Significance:** The physiological mechanisms underlying cold tolerance of insects are complex. Our study integrates tissue-specific responses with whole-body response to cold and compares these responses in different populations, phenotypes, and strains of the vinegar fly *Drosophila ananassae*. The study reveals that the response to extended cold in the renal and intestinal tissues are different, that the tissue-specific responses can be masked in the whole body, and that the transcriptomic response to extended cold can depend on population of origin, cold tolerance phenotype, or strain. Overall, this study sheds light on insect cold adaptation in terms of transcriptomic response to cold acclimation in the ionoregulatory tissues and whole bodies.

## Introduction

Temperature is a major factor influencing how species adapt to changing environments and colonize new ecosystems. How animals generally cope with thermal stressors and which genes are specifically responsible for cold adaptation has been a topic of intense research for decades in molecular and evolutionary biology. Recent developments in high-throughput sequencing methods allow insights into molecular mechanisms at the whole transcriptome and genome level, but integrating the complementary information of various experimental designs remains challenging. For example, studies on small insects often rely on either systemic or tissue-specific methodologies or single phenotypes, resulting in incomplete conclusions (Andersen *et al*., 2015; Hoedjes *et al*., 2024). Therefore, integrating different experimental designs and phenotypes to study transcriptional responses to cold is critical to uncovering mechanisms underlying physiological adaptation. Elucidating the mechanisms that underlie adaptation to changing environments in natural populations and integrating different approaches can help us understand which species prove to be successful in the future and have potential applications for conservation, agriculture, and pest management.

Many insects evolved and diversified in warm climates (Bale & Hayward, 2010). Nevertheless, insects have developed various strategies to cope with thermal stress and to successfully expand their thermal and habitat ranges (Overgaard & MacMillan, 2017). Some insects even tolerate freezing and remain active, such as the Antarctic midge *Belgica antarctica* (Teets *et al*., 2012) or the alpine beetle *Pytho deplanatus* (Ring, 1982). However, a wide range of insect species suffer from chilling injuries and eventually die when they are exposed to cold. This chill-susceptibility has been reported for several orders of insects (Overgaard & MacMillan, 2017). Chill-susceptible species include some of great economic and ecological importance, such as the honeybee *Apis mellifera* (Free & Spencer-Booth, 1960), but also agricultural pests, for instance, the migratory locust *Locusta migratoria* (Bayley *et al*., 2018) and disease vectors such as mosquitoes (Reinhold *et al*., 2018).

The physiology of cold stress resistance in insects is highly complex. When insects are exposed to cold, their ion and water homeostasis is disrupted (MacMillan *et al*., 2015). Additionally, transcriptomic and metabolic profiles are changed (MacMillan *et al*., 2016). Along with these systemic changes, chilling injuries affect the body at the cell level as they cause phase changes in the cell membrane, resulting in a stiff membrane with decreased permeability (Steponkus, 1984; Koštál, 2010; Drobnis *et al*., 1993). Further, cold exposure results in an imbalance of passive diffusion and active ion transport (MacMillan & Sinclair, 2011; Koštál *et al*., 2004; Zachariassen *et al*., 2004). An excess of potassium ions in the extracellular space leads to depolarization of the membrane (Findsen *et al*., 2014; Hoyle, 1953), eventually activating the voltage-gated calcium channels (Salkoff & Wyman, 1983; Findsen *et al*., 2016). The cellular influx of calcium ions through voltage-gated channels activates apoptotic and necrotic pathways, leading to chill injury (Koštál *et al*., 2004, 2007, 2016; Zachariassen *et al*., 2004; MacMillan *et al*., 2015, 2016). Consequently, the regulation of ion and water homeostasis has been identified as a key physiological feature for cold tolerance in insects with specifically the ionoregulatory tissues being essential for maintaining the water-ion balance in an insect body (Andersen *et al*., 2017; Andersen & Overgaard, 2020; Des Marteaux *et al*., 2018; Gerber & Overgaard, 2018; MacMillan *et al*., 2015, 2016, 2017; Yerushalmi *et al*., 2018). Despite several in-depth studies aiming to resolve this complexity, it remains challenging to identify and functionally characterize specific genes and genetic elements that contribute to the diversity of cold stress-resistant phenotypes (Udaka *et al*., 2010; von Heckel *et al*., 2016; MacMillan *et al*., 2016). With their worldwide distribution, Drosophilids are excellent models for studying these questions.

Flies from the genus *Drosophila* suffer from impaired synaptic transmission and muscular failure when they are exposed to cold; in other words, they lose their neuromuscular coordination and enter a comatose condition referred to as chill coma in which they are not able to move, feed, mate or escape predators (Kelty *et al*., 1996; MacMillan & Sinclair, 2011; Findsen *et al*., 2014). However, flies can recover from such a coma given that time and temperature of exposure to cold do not exceed certain thresholds. The time the fly needs for recovery from a cold-induced coma (chill coma recovery time, CCRT) is widely used to assess the cold hardiness of insects in the laboratory and serves as a predictor for the climatic condition at the geographical sampling site of a particular species (Gibert *et al*., 2001). Further, CCRT follows latitudinal clines within *Drosophila* species in Australia (Hoffmann *et al*., 2002; Hoffmann & Weeks, 2007; Hallas *et al*., 2002), Asia (Sisodia & Singh, 2010), Africa and Europe (David *et al*., 2003). Cold tolerance in Drosophilids is polygenic. Despite this complexity, variation in gene expression of only a few loci is associated with cold tolerance in subtropical populations of *Drosophila ananassae* (Königer & Grath, 2018), and variation of just three genomic regions explained more than 60% of the phenotypic variation between cold-tolerant and cold-sensitive strains (Königer *et al*., 2019). Recently, we provided Cas9-integrated fly strains that facilitate the use of the CRISPR-Cas9 system in this species and allow functional characterization of individual candidate loci (Yılmaz *et al*., 2023). In sum, *D. ananassae* has been established as a promising model for studying the genetic basis of cold tolerance.

Previous studies mostly neglected the potential of integrating two specific aspects to better inform about cold tolerance in insects. First, how insects respond to low temperatures can be plastic: the ability to tolerate cold and the genes responsible for the survival can change depending on prior exposure to cold. Such acclimation has been shown to alleviate the adverse effects of cold and can increase cold tolerance (Bubliy *et al*., 2002; Enriquez & Colinet, 2019; Hoffmann & Watson, 1993; MacMillan *et al*., 2016, 2017; Watson & Hoffmann, 1996; Yılmaz *et al*., 2023). Cold acclimation triggers substantial changes in the transcriptional response to cold in *Drosophila suzukii* (Enriquez & Colinet 2019). It reorganizes the transcriptome and metabolome of whole animals (MacMillan *et al*., 2016) and the gut transcriptome in *Drosophila melanogaster* (MacMillan *et al*., 2017). Second, conclusions on potentially ecologically relevant adaptations should not rely on a single phenotype (Andersen *et al*., 2015). Consequently, incorporating distinct cold tolerance phenotypes and systemic and ionoregulatory tissue-specific transcriptome data will allow pinpointing cold-related physiological adaptations.

In this study, we use isofemale lines collected from a population of ancestral origin in Bangkok, Thailand (from here on mentioned as ancestral population) and a derived population originating from Kathmandu, Nepal (Das *et al*., 2004). We examined the transcriptional response to cold acclimation in both the ancestral and the derived populations to shed light on the genetic mechanisms underlying cold tolerance. In addition, we integrated the original results of this study with previously gained insight into the distinct phenotype of cold shock. To this end, we contrasted the transcriptional response to cold at four levels: (1) population (ancestral vs. derived), (2) strain (cold-sensitive vs. cold-tolerant), (3) tissue (whole-body vs. ionoregulatory tissues, Malpighian tubules and hindgut), and (4) phenotype (cold acclimation vs. cold shock). We find empirical evidence for the influence of population and tissue on gene expression variation upon cold stress. Using an RNA sequencing approach that combined systemic with tissue-specific transcriptomes, we show that cold acclimation improves cold tolerance and results in more than 50% of the transcriptome being differentially expressed in the whole body and the ionoregulatory tissues.

Our results demonstrate that an RNA sequencing strategy combining a systemic overview with tissue-specific data can be a cost-effective and informative way to elucidate genes that have major effects on a crucial adaptation-related trait, namely, thermoregulation.

## Results

### Complementary elucidation of transcriptional response to cold acclimation with systemic whole body and tissue-specific data

Flies from ancestral and derived populations were previously described as having different sensitivity to cold shock as determined by chill coma recovery time, CCRT (Königer & Grath, 2018). Additionally, preliminary tests revealed that acclimation to cold improves the tolerance and decreases the chill coma recovery time (see Supplementary Text). To receive an informative overview of the transcriptional response to cold acclimation in *D. ananassae*, we combined high-throughput RNA sequencing of the whole body (wb), as well as of two ionoregulatory tissues, Malpighian tubules (Mt) and hindguts (hg). We obtained RNA sequencing data from male flies upon 4-day acclimation at 10°C. In the whole body and hindgut samples, 54% of all expressed genes were differentially expressed in response to cold acclimation, and the percentage of DE genes in the Malpighian tubules was 50% (Table 1). The principal component analysis of all samples revealed that they clustered according to the tissue RNA was extracted from, whereas no separation according to the treatment was observed (Figure 1A). The first principal component explained 68% of the variance, separating the ionoregulatory tissues from the whole-body samples. The second principal component explained 19% of the variance, separating the two ionoregulatory tissues. Though pairwise comparisons revealed that 2952 differentially expressed genes were shared between sample types, a considerable number of genes (ranging from 505 genes that were explicitly up-regulated in Malpighian tubules to 1436 genes that were down-regulated explicitly in the whole body) were specifically up- or down-regulated only in the individual tissues (Figure 1B-C). Notably, these genes were enriched in distinct GO categories (Supplementary Table 3-4). These results suggest that the combination of systemic and informed tissue-specific RNA sequencing data allows for complementary conclusions on cold tolerance in *Drosophila*.

**Figure 1:**
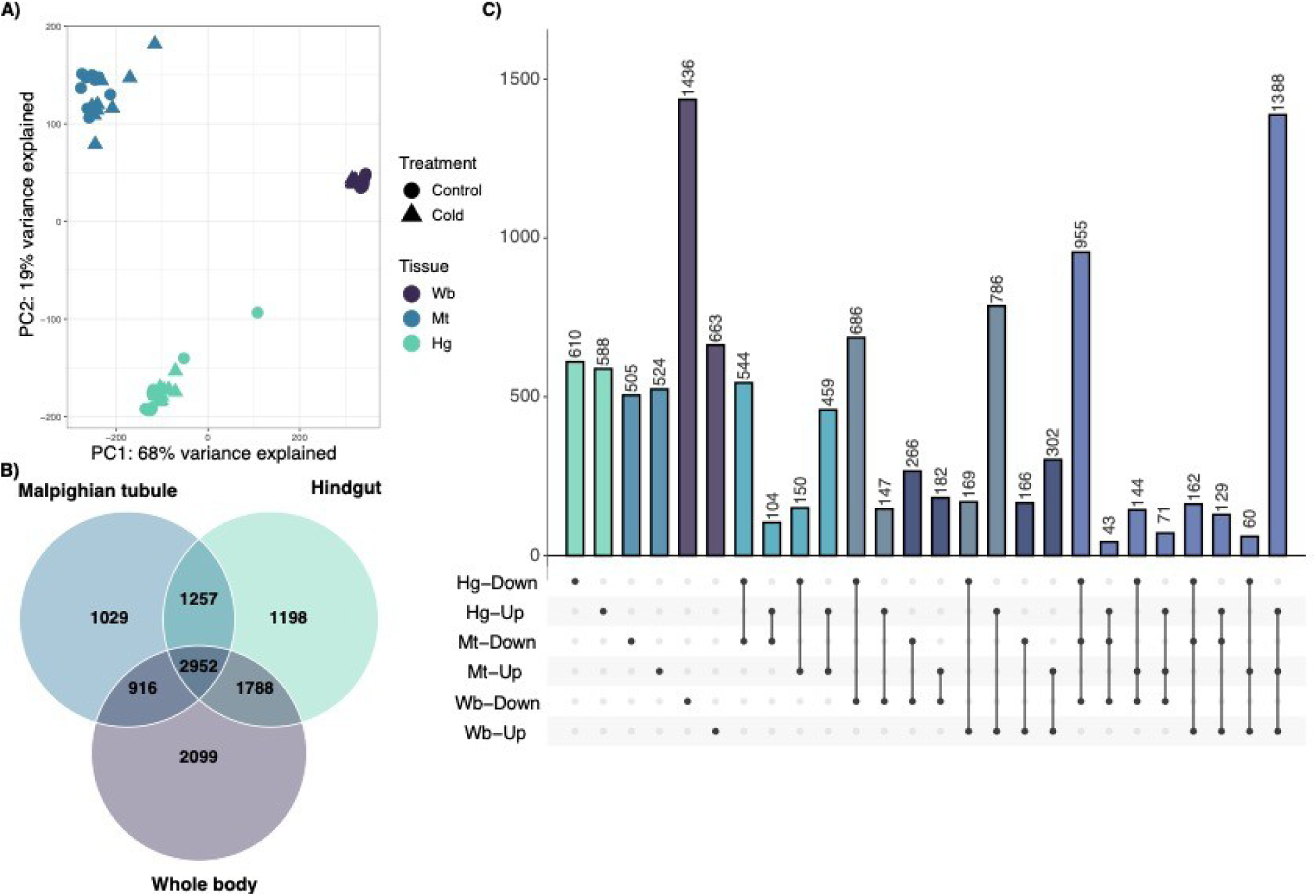
Transcriptomic response to cold acclimation in ionoregulatory tissues and the whole body. A) Principal component analysis. The samples are shown according to the tissue (color) and treatment (shape). Whole body (Wb), Malpighian tubule (Mt), and hindgut (Hg) tissues are represented by purple, blue, and green colors, respectively. The control samples are represented by circles and the cold acclimated samples are represented by triangles. B) Venn diagram showing the number of DE genes specific to or shared among sample types: Whole body, Malpighian tubule and Hindgut samples exhibited both shared, as well as exclusive differentially expressed genes. C) Upset plot showing differentially expressed genes and their overlap between the three different sample types. Black, horizontal bars indicate the total number of differentially expressed genes that were significantly up- or down-regulated in a sample in response to cold acclimation. A black circle indicates that a sample and direction (up/down) combination is included in an intersection class. Intersection classes with a single circle are comprised of genes significantly differentially expressed in a single tissue and direction. Circles connected by a line indicate intersection classes comprised of multiple tissue and direction combinations. Vertical bars correspond to the number of genes in each intersection class, colors represent the respective sections of the Venn diagram.

**Table 1:** Number of differentially expressed genes in different sample types regarding the general, population-specific, phenotype-specific, and line-specific response to cold acclimation.

|  | total expressed<br>genes | up<br>regulated | down regulated | total DE |
| --- | --- | --- | --- | --- |
| **General response to acclimation** |  |  |  |  |
| Whole body | 14304 | 3825 (27%) | 3930 (27%) | 7755 (54%) |
| Malpighian tubule | 12189 | 3280 (27%) | 2874 (24%) | 6154 (50%) |
| Hindgut | 13272 | 3715 (28%) | 3480 (26%) | 7195 (54%) |
| **Population-specific response** |  |  |  |  |
| <u>Whole body</u> |  |  |  |  |
| BKK | 14300 | 3618 (25%) | 3668 (26%) | 7286 (51%) |
| KATH | 14300 | 3414 (24%) | 3497 (24%) | 6911 (48%) |
| KATH vs BKK* | 13191 | 1181 (9%) | 827 (6.3%) | 2008 (15%) |
| <u>Malpighian tubule</u> |  |  |  |  |
| BKK | 11721 | 3077 (26%) | 2606 (22%) | 5683 (48%) |
| KATH | 11191 | 2288 (20%) | 2174 (19%) | 4462 (40%) |
| KATH vs BKK* | 9125 | 124 (1.4%) | 92 (1%) | 216 (2.4%) |
| **Phenotype-specific response** |  |  |  |  |
| <u>Hindgut</u> |  |  |  |  |
| Fast | 12821 | 3203 (25%) | 3570 (28%) | 6773 (53%) |
| Slow | 12547 | 2907 (23%) | 2784 (22%) | 5691 (45%) |
| Fast vs Slow* | 12547 | 182 (1.5%) | 246 (2%) | 428 (3.4 %) |
| **Line-specific response\*\*** |  |  |  |  |
| <u>Whole body</u> |  |  |  |  |
| FastBKK | 14300 | 3333 (23%) | 3434 (24%) | 6767 (47%) |
| SlowBKK | 14300 | 3091 (22%) | 3300 (23%) | 6391 (45%) |
| FastBKK vs SlowBKK* | 13468 | 508 (3.8%) | 472 (3.5%) | 980 (7.3%) |

\* For these comparisons, the columns show the number of genes that are significantly differentially expressed between the compared groups. Up and down-regulated columns indicate direction regarding the effect of acclimation on the former group compared to the latter group. BKK = Bangkok, KATH = Kathmandu.
\*\* Only isofemale lines of the ancestral population (BKK) are taken into consideration. The terms 'fast' and 'slow' indicate faster and slower recovery, respectively.

### Strong systemic transcriptional response to cold acclimation

We performed RNA sequencing on whole-body samples to obtain a systemic overview of the transcriptional response to cold acclimation in *Drosophila ananassae*. This approach has the unique advantage of being both cost-efficient and unbiased in terms of preconditions on transcription. Samples of acclimation and control treatments were clustered according to the populations of origin (Figure 2A). The first principal component explained 58% of the variance. It separated cold-acclimated samples from control samples, while the second principal component explained 14% of the variance and separated the derived population from the ancestral population. Within the ancestral population, the samples also clustered according to their CCRT phenotype (Figure 2B). We addressed three questions with this data: (1) What is the general, systemic transcriptional response to cold acclimation? (2) How does transcriptional response differ between ancestral and derived populations? (3) How does transcriptional response differ between cold-tolerant and cold-sensitive strains in the ancestral population?

**Figure 2:**
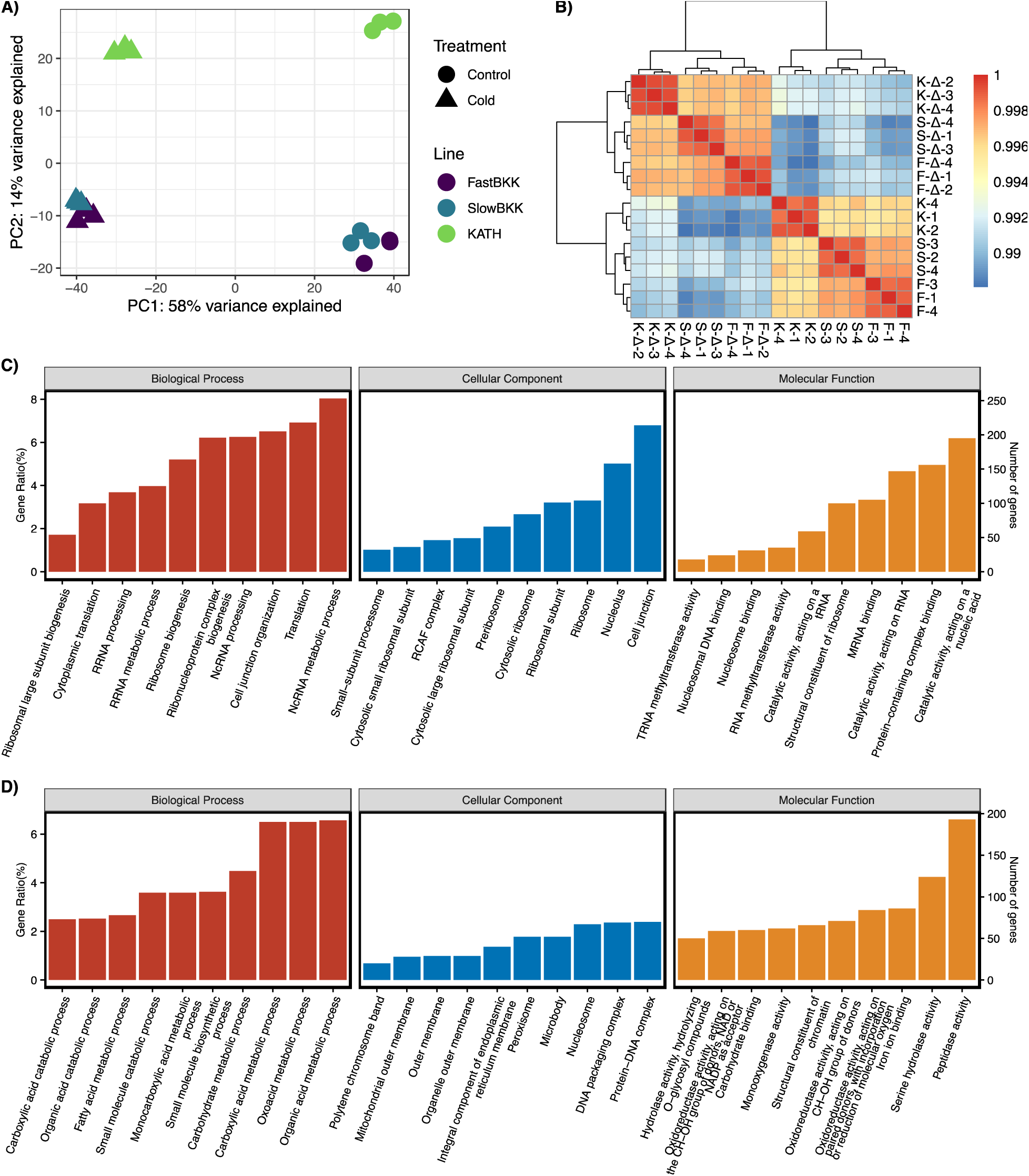
Transcriptomic response to cold acclimation in the whole body. A) Principal component analysis. The samples are shown according to the pooled strains (color) and treatment (shape). FastBKK, SlowBKK, and KATH sample pools are represented by purple, blue, and green colors, respectively. The control samples are represented by circles and the cold acclimated samples are represented by triangles. B) Heatmap for the sample pools. Sample clusters are shown on the left and top of the heatmap. The letters indicate the sample pools FastBKK (F), SlowBKK (S), Kath (K). Cold acclimated samples are indicated by delta (Δ) sign. The numbers correspond to the replicates. The similarity matrix indicates high (red) and low (blue) similarity of samples. Gene ontology enrichment analysis results for genes C) up-regulated and D) down-regulated in the whole body in response to cold acclimation. Biological processes (red bars), cellular components (blue bars), and molecular functions (yellow bars) are shown in separate panels.

More than half of the transcriptome (54%) was differentially expressed between control and treatment, with 3825 genes significantly up-regulated and 3930 genes significantly down-regulated (Table 1). Gene ontology enrichment analysis revealed that up-regulated genes were enriched in biological processes related to translation and cell junction organization (Figure 2C, Supplementary Table 5). On the contrary, down-regulated genes were enriched in metabolic processes (Figure 2D, Supplementary Table 5). These results suggest a solid response to cold acclimation in the whole body, and the transcriptional response of populations differs not only in the presence of cold treatment but also at the basal level.

### Systemic differences between populations are mediated by genes involved in detoxification and maintaining ion balance

To characterize the differences between populations, we contrasted the transcriptional response of ancestral and derived populations. A total of 14300 genes were expressed in each population (Table 1). In the ancestral population, 51% of these genes were differentially expressed upon cold acclimation, while 3618 genes were significantly up- and 3668 down-regulated, respectively. In comparison, 48% of these genes were differentially expressed in the derived population, while 3414 genes were significantly up-regulated, and 3497 genes were significantly down-regulated. Although the populations shared over half of the differentially expressed genes (Supplementary Figure 2A), 2008 genes showed significant differential expression between populations in response to cold acclimation. The GO analysis suggested that the genes with differential expression between the populations were mainly located in muscle tissues and enriched in cilium movement and various metabolic processes, including fatty acid metabolism (Supplementary Figure 2B). Notably, these genes were also enriched in molecular functions related to ion binding, ion channel, and oxidoreductase activity (Supplementary Table 6). Overall, these results suggest the difference between the acclimation response of populations ground on ion transport and detoxification.

### The actin cytoskeleton differentiates the acclimation response of tolerant and sensitive strains in the ancestral population

To address the question of how the transcriptional response differs within the ancestral population between cold-tolerant and cold-sensitive strains, we compared gene expression upon cold acclimation in both types of strains. For the cold-tolerant strains, 3333 genes were significantly up-regulated, and 3434 genes were significantly down-regulated (Table 1). For the sensitive strains, 3091 genes were significantly up-regulated, and 3300 genes were significantly down-regulated. However, 980 genes were differentially expressed between cold-sensitive and cold-tolerant strains. GO enrichment analysis revealed significant enrichment of these genes in various biological processes related to the actin cytoskeleton. (Supplementary Table 7)

### Adaptive transcriptional response is evident in the renal tissues

Malpighian tubules (Mt) are the renal equivalent of mammals in insects and contribute to maintaining homeostasis, which can be disrupted by cold exposure. Therefore, we investigated the transcriptional response of Mt to characterize the impact of cold acclimation on renal tissues in *D. ananassae*. Like the whole-body samples, RNA samples extracted from Mt of acclimated and control flies were clustered according to the populations of origin, suggesting that the populations are distinct in their response to acclimation and at the basal level of gene expression (Figure 3A). Principal component analysis indicated that 37% of the variation can be explained by the first principal component, which separated cold-acclimated and control samples, and 14% of the variation can be explained by the second principal component, which separated the ancestral and derived populations. Therefore, we investigated the general transcriptional response of Mt to cold acclimation and the population-specific response.

**Figure 3:**
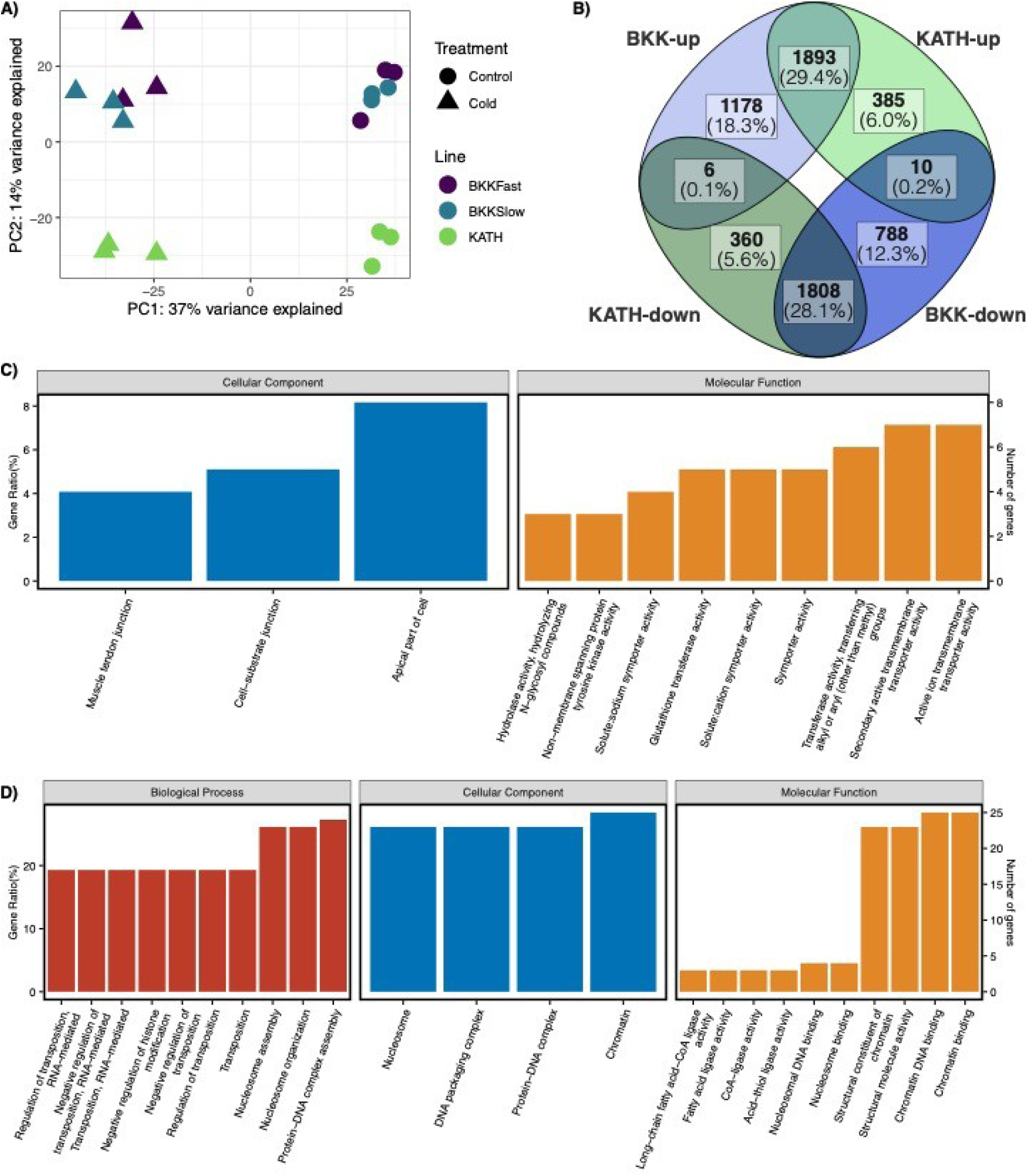
Transcriptomic response to cold acclimation in the Malpighian tubules. A) Principal component analysis. The samples are shown according to the pooled strains (color) and treatment (shape). FastBKK, SlowBKK, and KATH sample pools are represented by purple, blue, and green colors, respectively. The control samples are represented by circles and the cold acclimated samples are represented by triangles. B) Venn diagram showing the number of DE genes specific to or shared between populations: Bangkok and Kathmandu populations exhibited both shared, as well as exclusive differentially expressed genes in response to acclimation. Gene ontology (GO) enrichment analysis results for genes C) up-regulated and D) down-regulated in the Malpighian tubules of Kathmandu population compared to Bangkok population in response to cold acclimation. Biological processes (red bars), cellular components (blue bars), and molecular functions (yellow bars) are shown in separate panels.

In the Mt samples, 3280 genes were up-regulated, and 2874 were down-regulated (Table 1). Similar to the whole-body samples, up-regulated genes of Mt were enriched in GO terms related to translation and transcription (Supplementary Figure 3A, Supplementary Table 8), and the down-regulated genes were enriched in processes related to fatty acid degradation (Supplementary Figure 3B, Supplementary Table 8). However, contrary to the whole-body samples, the down-regulated genes of the Mt were also enriched in chromatin remodeling. Additionally, a major part of the down-regulated genes was located in the mitochondrion and, more specifically, the respiratory chain complex.

### Differences in renal tissues of populations are mediated by genes involved in ion transport and lipid metabolism

In order to understand the differences in metabolic processes involved in cold adaptation between the ancestral and derived populations, we compared the transcriptional response to cold acclimation in flies from the Bangkok and Kathmandu populations. In the ancestral population, 11721 genes were expressed, with 3077 up-regulated and 2606 down-regulated (Figure 3B, Table 1). In the derived population, 11191 genes were expressed, with 2288 up-regulated and 2174 down-regulated. Among these DE genes, 216 genes showed differential expression between the populations. We conducted a GO analysis to explore the functions of these genes and found GO terms related to transposition. To enhance the analysis, we performed separate analyses for gene sets: genes up regulated in the derived population compared to the ancestral one, and *vice versa*. The analysis showed that genes up regulated in the Kathmandu population compared to Bangkok were enriched in molecular functions related to ion transport (Figure 3C, Supplementary Table 9). Conversely, genes up regulated in the ancestral population compared to the derived one were enriched in processes and functions related to transposition (Figure 3D, Supplementary Table 9). Additionally, this analysis revealed an enrichment in molecular functions related to fatty acid ligase activity.

### Hindgut responds to cold acclimation with respect to phenotype, but not to population

Along with the Malpighian tubules, hindguts (hg) are essential in regulating ion and water balance. Therefore, we investigated the tissue-specific transcriptional response to cold acclimation in the hindgut of *D. ananassae* flies to elucidate molecular mechanisms of cold adaptation in insects. Initial PCA revealed two clusters and an outlier sample (Figure 4A). When the outlier was removed to increase the resolution, PC1 explained 48% of the variance, and PC2 explained 12%, separating the treatments and phenotypes, respectively (Supplementary Figure 4A). The acclimated samples clustered according to their cold tolerance phenotype, in other words, according to their recovery time from chill coma. However, as opposed to the whole body and MT, the control samples showed no apparent separation, which suggests that the strains do not differ at their basal gene expression level according to their cold tolerance phenotype or population of origin. Therefore, we addressed two questions with this data: (1) What is the general response to cold acclimation in the hindguts? (2) How does transcriptional response differ between cold-tolerant and cold-sensitive strains?

**Figure 4:**
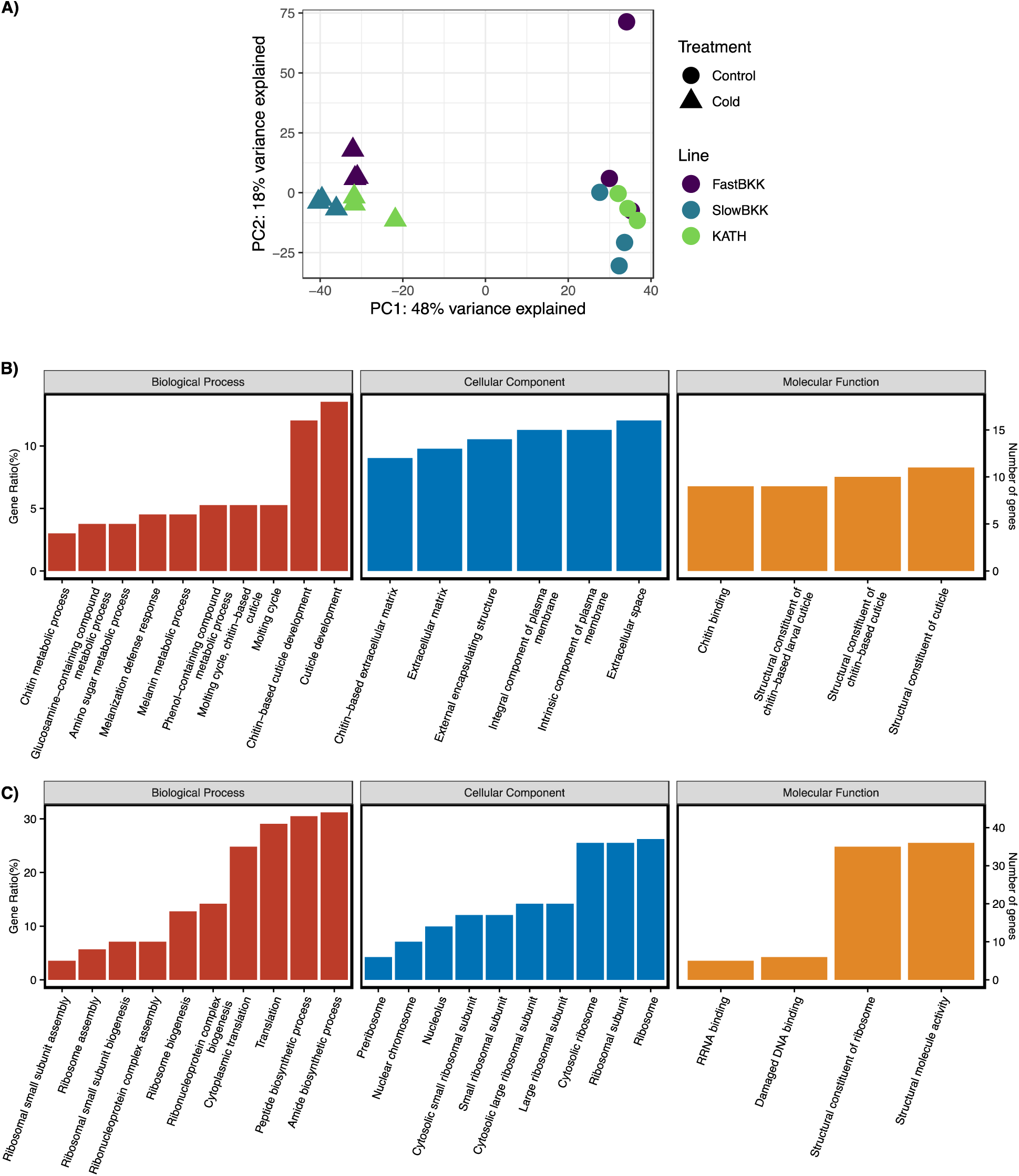
Transcriptomic response to cold acclimation in the hindguts. A) Principal component analysis. The samples are shown according to the pooled strains (color) and treatment (shape). FastBKK, SlowBKK, and KATH sample pools are represented by purple, blue, and green colors, respectively. The control samples are represented by circles and the cold acclimated samples are represented by triangles. Gene ontology (GO) enrichment analysis results for genes B) up-regulated and C) down-regulated in the hindguts of tolerant flies (fast phenotype) compared to sensitive flies (slow phenotype) in response to cold acclimation. Biological processes (red bars), cellular components (blue bars), and molecular functions (yellow bars) are shown in separate panels.

In the hindgut samples, we observed a significant up-regulation of 3835 genes and a down-regulation of 3483 genes in response to cold acclimation (Table 1). Like the whole body and Malpighian tubules, the up-regulated genes in the hindguts were enriched in GO terms related to translation and transcription (Supplementary Figure 4B, Supplementary Table 10). In contrast, the down-regulated genes showed enrichment in biological processes related to cellular respiration and lipid metabolism, with many of these genes located in the mitochondrial membrane (Supplementary Figure 4C, Supplementary Table 10).

### Cellular respiration and chitin metabolism play roles in the acclimation response of tolerant and sensitive strains in the hindgut

We characterized the transcriptional response to cold acclimation in cold-tolerant and cold-sensitive strains with differential gene expression analysis. In the cold-sensitive strains, 3046 genes were significantly up-regulated, and 2886 genes were significantly down-regulated in response to acclimation (Table 1). On the other hand, in the cold-tolerant strains, 3401 genes were significantly up-regulated, and 3224 genes were significantly down-regulated in response to cold acclimation. Between the tolerant and sensitive strains, 330 genes were differentially expressed. GO enrichment analysis revealed that these genes were enriched in biological processes related to translation and ribosome biogenesis and in molecular functions related to chitin binding. We separately investigated the genes up-regulated in the cold-tolerant strains compared to the sensitive strains and *vice versa*. Genes significantly up-regulated in the cold-tolerant strains compared to the cold-sensitive strains were enriched in GO terms related to cuticle development, respiration, and metabolism (Figure 4B, Supplementary Table 11). Additionally, the products of these genes were localized mainly to the membrane and extracellular matrix. Genes significantly up-regulated in the cold-sensitive strains compared to the cold-tolerant strains were enriched in GO terms related to translation (Figure 4C, Supplementary Table 11).

### Distinct transcriptional response to cold acclimation and cold shock

To receive a comprehensive overview of cold tolerance measured by distinct phenotypes, we contrasted our previous results of the transcriptional response to cold shock as measured by chill coma recovery time upon cold shock (Königer & Grath, 2018) with the transcriptome data from this study. To this end, we considered the genes differentially expressed in the cold-tolerant and cold-sensitive strains of the ancestral population from Bangkok, Thailand, in two conditions: (1) cold shock and (2) cold acclimation.

Our findings reveal the intricate nature of gene expression in response to cold shock and cold acclimation. We observed that more genes are differentially expressed in response to cold acclimation than cold shock (Supplementary Figure 5), indicating a more complex regulatory network in the former. To understand these responses, we investigated the up- and down-regulated genes in both treatments and performed a GO enrichment analysis.

The genes up-regulated in response to both treatments were enriched in actin cytoskeleton-related GO terms (Supplementary Table 12). In contrast, the down-regulated ones showed enrichment in various metabolic processes such as organic acid, lipid, and carbohydrate metabolic processes (Supplementary Table 12). We also investigated the genes with opposite expression (i.e., up-regulated in one treatment but down-regulated in the other) in response to treatments. Among the 1932 genes that showed differential expression in both studies, 390 genes had opposite directions. The genes up-regulated upon cold shock but down-regulated upon acclimation were enriched in the immune response (Supplementary Table 13). On the other hand, the genes that were up-regulated in response to acclimation but down-regulated after cold shock showed enrichment in biological processes related to transcription and translation (Supplementary Table 13).

Finally, we investigated the genes that show differential expression in response to only one of the treatments. Over 5000 genes were up- or down-regulated upon acclimation, but only 1070 genes were uniquely expressed after cold shock. The genes differentially expressed only upon the cold shock treatment were enriched in protein folding, actin cytoskeleton organization, replication, and glycosylation (Supplementary Table 14). The genes that were differentially expressed upon acclimation showed enrichment in translation and transcription as well as amino acid metabolism and copulation (Supplementary Table 14).

## Discussion

Understanding the differences between the ancestral and derived populations and the variance within the ancestral Bangkok population sheds light on the adaptive processes leading to increased cold tolerance of the species. The two populations were tested for their acclimation response. Both populations showed improved cold tolerance upon acclimating to mild temperatures as expected, but the underlying mechanisms were unknown. Comparative transcriptomics using the whole-body samples provided the systemic overview to understand the differences in the acclimation response between the two populations and within the ancestral population. Additionally, the tissue-specific response to acclimation provided further insight into population and phenotypic differences.

In a complex phenotype, such as cold response, transcriptomic analysis of a whole body might lead to missing information (reviewed in Hoedjes *et al*., 2024). However, tissue-specific RNA-Seq can unmask details otherwise lost in the systemic approach. Both whole-body and tissue-specific approaches were used in this study to increase the strength of the comparison of populations, phenotypes, and strains. The DE genes in the Malpighian tubule samples enriched in hypoxia response and HSP binding activity are not differentially expressed in the whole-body samples. Similarly, the DE genes in the hindgut samples enriched in sodium channel activity and cuticle structure are not differentially expressed in the whole body. On the other hand, even though the genes enriched in cellular respiration were up-regulated in the whole body, they were down-regulated in both ionoregulatory tissues. Taken together, combining whole-body and tissue-specific RNA-Seq allowed combining advantages while eliminating the disadvantages of each approach.

Previous studies on the cold acclimation response of other *Drosophila* species suggest that the percentage of differentially expressed genes in response to acclimation decreases as the species becomes more tolerant to cold (Parker *et al*., 2015, 2021; MacMillan *et al*., 2016; Enriquez & Colinet, 2019). Even though these differences can be due to technological advancements, acclimation conditions, or analysis methods, the current study’s large-scale differential expression aligns with this claim. In addition, the general response to cold acclimation consisted of differential expression of genes related to translation, transcription, and metabolism. Over-expression of translation-related genes parallels the high number of DE genes responding to acclimation. The populations also showed a slight difference in DE gene percentage, supporting the previous suggestion.

Exposure to low temperatures decreases membrane fluidity (Steponkus, 1984; Drobnis et al., 1993; Koštál, 2010). Homeoviscious adaptation suggests that the organism can adjust its membrane fluidity by increasing the proportion of unsaturated fatty acids or the ratio of short-chain fatty acids to long-chain fatty acids (Sinensky, 1974; Koštál, 2010). Insects can adjust their membrane composition through fatty acid metabolism to keep the membrane fluidity consistent, which may increase cold tolerance during acclimation (reviewed in Teets & Denlinger, 2013). Aquaporins, essential membrane proteins supporting cold tolerance in many insects (Zhao *et al*., 2023), were suggested to regulate cold shock response by modulating water balance and fatty acid metabolism in ticks (Wang *et al*., 2025). In our study, the fatty acid elongation was down-regulated in response to acclimation in both populations, preventing a decrease in the short-chain-to-long-chain ratio. However, the differential expression analysis between the populations suggested that the down-regulation of genes related to fatty acid elongase activity was significantly higher in the Bangkok population than in the Kathmandu population. These results suggest that the ancestral population prevents the elongation of short-chain fatty acids into long-chain ones to maintain the membrane fluidity. In contrast, the acclimation response of the derived population does not solely depend on membrane structure.

As discussed previously, cold exposure increases potassium ion concentration in the extracellular space, resulting in membrane depolarization (Hoyle, 1953; Findsen *et al*., 2014; reviewed in Overgaard & MacMillan, 2017 and in Overgaard *et al*., 2021). As acclimation increases tolerance, adjustments are expected to be made to improve the maintenance of the water-ion balance. Hence, the up-regulation of genes regulating ion transport mechanisms can improve the cold tolerance phenotype. The genes significantly up-regulated in the derived population compared to the ancestral one showed enrichment in ion transport and channel activity. Specifically, the up-regulation of genes regulating the potassium channel activity in the Kathmandu population indicates that the derived population takes advantage of preventing depolarization by maintaining the potassium concentration. On the other hand, in the Malpighian tubules, the ion transport genes were responsible for the active transmembrane transport rather than ion-specific channels. As active transport is affected more than passive diffusion in the case of cold exposure (Koštál *et al*., 2004; Zachariassen *et al*., 2004; MacMillan & Sinclair, 2011), maintaining the ion balance in response to acclimation depends on regulating the active transport mechanism.

Taken together, these results suggest that the ancestral population maintains the membrane fluidity to prevent the loss of water-ion balance, whereas the derived population can maintain the active transport in the Malpighian tubules and regulate the potassium ion concentration in the extracellular space in response to acclimation, improving the cold tolerance. Therefore, maintaining ion transport plays a crucial role in the adaptation of the derived Kathmandu population to cold temperatures.

Considering the phenotypic variance within and between populations, comparative transcriptomic analysis of the cold acclimation response also sheds light on the differences between the phenotypes. The analysis focused on the hindgut tissue, where cellular respiration and lipid metabolism-related genes were down-regulated in response to acclimation, whereas cuticle development was up-regulated.

A previous study showed that prolonged exposure to cold reduces insect respiration rate (Colinet *et al*., 2017). However, the same study also suggested that high tolerance can be achieved by maintaining the energy requirement under cold stress. In line with these suggestions, even though the respiration-related genes were down-regulated in response to acclimation in both phenotypes, the difference between the tolerant and sensitive phenotypes was significant: sensitive lines showed significant down-regulation of these genes compared to the tolerant lines. These indications suggest that maintaining the respiration-related processes can be associated with fast recovery from chill coma.

Together with respiration, lipid metabolism-related genes were significantly down-regulated in hindguts in response to cold acclimation. As discussed above, low temperatures affect the lipid bilayer structure of the membrane, and these impacts can be overcome by homeoviscious adaptation. Even though the genes associated with fatty acid elongase activity were down-regulated, no phenotype-specific difference was observed.

Previous studies showed that reabsorption in the hindgut plays a crucial role in cold tolerance of *Drosophila* species (Andersen *et al*., 2017; Andersen & Overgaard, 2020). Therefore, any adaptation preventing or minimizing the leak in the hindgut is expected to increase tolerance to cold. As the cuticle structure of the hindgut prevents water loss (Phillips, 1970), activation of related pathways is expected to be observed in cold-tolerant strains. Interestingly, genes related to cuticle development and chitin metabolism were up-regulated in the hindguts of all strains in response to cold acclimation. However, the up-regulation significantly differed between the phenotypes. As expected, tolerant strains showed more up-regulation of related genes than the sensitive strains, suggesting that cuticle formation plays a vital role in the cold tolerance level of the species.

The comparative transcriptomic analysis of the whole-body cold acclimation response also explains the variation within the ancestral population. Differences were observed in GO terms related to actin cytoskeleton and calcium channels between the fast and slow-recovering ancestral strains. Genes related to both were up-regulated in response to acclimation, but the upregulation was significantly more in the tolerant fast-recovering strains.

Actin cytoskeleton plays roles in various biological processes, including cell structure, shape, migration, and signaling. Previous studies suggested that the actin cytoskeleton is essential in cold tolerance (Cottam *et al*., 2006; Kim *et al*., 2006; von Heckel *et al*., 2016; Chen *et al*., 2017; Des Marteaux *et al*., 2018; Bowman *et al*., 2018). One of these studies also showed that the tolerant and sensitive Bangkok strains differed in actin polymerization: six genes had differential expression in response to cold shock (Königer & Grath, 2018). On the other hand, five out of six genes were up-regulated in both phenotypes in response to acclimation. However, they did not show phenotype-specific differential expression even though the initial expression levels were significantly lower in the sensitive phenotype.

In addition to the roles above, the actin cytoskeleton also acts as an anchor to ion channels and regulates ion transport (reviewed in Denker & Barber, 2002). Previous studies showed that together with microtubules, actin cytoskeleton dynamics affect the calcium ion influx. Calcium ion concentration is shown to be affected by cold temperatures (Salkoff & Wyman, 1983; Findsen *et al*., 2016) and plays an essential role in the activation of apoptotic and necrotic pathways (Koštál et al., 2004, 2007, 2016; Zachariassen et al., 2004; MacMillan et al., 2015, 2016). Genes related to calcium channels were also up-regulated in response to acclimation in the ancestral population. However, the differential expression analysis between the two phenotypes of the ancestral population showed that the calcium transport-related genes were significantly more up-regulated in the fast-recovering phenotype than in the slow-recovering one. These results suggest that the acclimation response of the two phenotypically distinct ancestral pools differ in actin cytoskeleton and ion transport, focusing on calcium ions.

In this study, we showed that acclimation to mild temperatures profoundly affected the cold tolerance of *D. ananassae* and investigated the mechanisms leading to increased tolerance. We showed that different populations had distinct transcriptomic responses to acclimation in the whole body and the Malpighian tubules. The differences between populations manifested through ion transport and lipid metabolism. In addition, the cold tolerance phenotype affected the acclimation response in the hindgut: tolerant strains could better maintain the cellular respiration that may generate the energy required by the active transport mechanism and took advantage of the cuticle structure that prevents water loss. Finally, we showed that the phenotypic variation within the ancestral population resulted in differences in the acclimation response, including actin cytoskeleton and calcium channel-related processes.

Further studies should be performed to validate these differences between the two phenotypes, focusing on the function of the hindgut and cellular respiration. Mitochondria activity has been described to undergo various changes under cold stess in insects, both to minimize cell damage as well as due to a decline in mitochondrial homeostasis caused by cell injuries and also depending on experimental conditions, what should be investigated further (Lubawy *et al*., 2022). Hindgut reabsorption functional assays were previously used in larger insects such as grasshoppers (Gerber *et al*., 2021) but also successfully performed in other *Drosophila* species (Andersen & Overgaard, 2020; Andersen *et al*., 2017). Similarly, cellular respiration measurements were successful in other species (Colinet *et al*., 2017) and can be used in *D. ananassae*. In addition to functional assays, candidate genes arising from further analyses can be edited through CRISPR-*Cas9* technology previously established in the species (Yılmaz *et al*., 2023) for functional validation.

## Materials and Methods

### Fly strains

This study used eight *D. ananassae* isofemale lines from Thailand (Bangkok, BKK) and three isofemale lines from Nepal (Kathmandu, KATH). The lines from Bangkok were the same as used in previous studies (Königer and Grath, 2018; Königer *et al*., 2019). All lines were collected in 2002 (Das *et al*., 2004). The fly strains were maintained under standard laboratory conditions (Rearing temperature – RT: 22°C ± 1°C 14:10 hours light:dark cycle) at low density on corn-molasses medium. Experiments were performed on flies that were expanded in large vials for two generations under the same conditions.

### Cold Acclimation Assay

Acclimation conditions were determined via preliminary tests (see Supplementary Text, Supplementary Methods). Acclimation vials contained standard food and were kept at 10°C for 24 hours prior to the experiment to allow the food to cool down to acclimation temperature. Male flies were pooled per population and phenotype on the day of emergence. After 24 hours, flies were transferred to vials containing standard medium as described above. The vials were kept at 10°C for 4 days, followed by 90 minutes recovery time at rearing temperature.

### RNA Extraction and Sequencing

Ten individual males per sample were snap-frozen in liquid nitrogen for whole-body samples, and RNA was extracted using MasterPure™ RNA Purification Kit (Epicentre, Madison, WI, USA) following the manufacturer’s protocol. For hindgut and Malpighian tubule samples, tissues from 60 males per sample were dissected, and the tissues were collected into 1.5 mL Eppendorf tubes containing 10-20 µl 1X PBS. RNA extraction from tissues was performed using RNeasy Mini Kit (Qiagen). RNA quantity and quality for all samples were confirmed with NanoDrop© (ND 1000, VWR International, Radnor, PA, USA) and a bioanalyzer (Bioanalyzer 2001, Agilent Technologies, Santa Clara, CA, USA, provided by the LMU genomics service unit). All samples were sent for sequencing to an external facility (Novogene, Cambridge, UK), which performed poly-A mRNA purification, reverse transcription, and high-throughput using a Novaseq 6000 sequencer (Illumina, San Diego, CA, USA) to generate 150-bp paired-end reads. On average, 88 million reads were generated per sample. An average of 73 million reads (83%) were successfully mapped to the *D. ananassae* transcriptome (dana_r1.05_FB2016_01) using NextGenMap (version 0.5.2) (Sedlazeck *et al*., 2013) and the annotation of FlyBase release 1.05. This release was used to compare the results of the present study with previously described data on transcriptome response following cold shock (Königer and Grath, 2018). All subsequent analyses were performed in R (version 4.2.2) (R Core Team, 2022).

### Differential Gene Expression Analysis

Differentially expressed genes were called using DESeq2 (version 1.38.3) (Love *et al*., 2014). The general response to treatment was assessed using ∼Treatment as the design formula. The differences between other variables (populations, phenotypes, or strains) were assessed using the design formula ∼Variable + Treatment + Variable:Treatment. Significant log2 fold changes between control and cold-acclimated samples were determined using a Wald test and a subsequent p-value multiple testing correction according to Benjamini-Hochberg. Genes with a corrected p-value less than 0.05 were considered as significantly differentially expressed.

### Gene Ontology (GO) Enrichment Analysis

GO analysis was performed using clusterProfiler (version 4.6.2) (Wu *et al*., 2021) implemented in R to determine the GO terms associated with *D. melanogaster* orthologs of the differentially expressed genes. The p-values were corrected using FDR method with a p-value cut-off of 0.01. The significant GO terms were visualized using the genekitr package (version 1.2.5) (Liu, 2023).

### Comparison with Cold Shock

Previously, Königer and Grath (2018) analyzed differential gene expression before, at 15 minutes, and 90 minutes after a cold shock in cold-sensitive and cold-tolerant strains of the Bangkok population. The study used the same Bangkok strains used in the current study, but the samples were not pooled for RNA extraction. Additionally, the data contained an extra time point (15 minutes after cold shock), which we did not test in the current study. Therefore, we used mapped cold shock reads that were kindly provided by Königer and Grath, analyzed them with the same DESeq2 model as the cold acclimation reads, and compared the DE genes between two phenotypes of the BKK population upon cold acclimation to the DE genes between the same groups at 90 minutes after cold shock.

## Data Availability

Gene expression data are available at GEO under the accession number GSE270239. All other data and scripts are available in the article and in its Supplementary Material.

## Author Contributions

Conceptualization S.G.; investigation V.M.Y., Z.B.; writing-original draft V.M.Y., S.G.; visualization V.M.Y., S.G.; funding acquisition S.G.

## Acknowledgements

Our work was supported by the Deutsche Forschungsgemeinschaft (project 271330745 to S. G.). We thank Hilde Lainer and Bianca Hölldobler for technical assistance in the laboratory, Amanda Glaser-Schmitt and John Parsch for their valuable input on the analysis of raw reads, and Annabella Königer for cold shock read counts. We extend our appreciation to all current and previous members of the Division of Evolutionary Biology, as well as the Graduate School Life Science Munich (LSM) at Ludwig-Maximilians-University Munich (LMU).

