## Supplementary Text and Methods for "Navigating the Cold: Integrative Transcriptome Sequencing Approach Reveals Ionoregulatory and Whole-Body Responses to Cold Acclimation in *Drosophila ananassae*"

#### Chill Coma Recovery Time Improves with Acclimation to Mild Temperatures

Prior exposure to mild stress conditions can improve the tolerance phenotype. To test the effect of temperature and duration of acclimation on CCRT, we tested four temperatures (6-12˚C) for different periods (1-5 days). We used three strains representing fast-recovering Bangkok (BKK12 for FastBKK), slow-recovering Bangkok (BKK13 for SlowBKK), and Kathmandu (KATH14 for KATH) populations. Flies exposed to 6˚C for more than a day showed high mortality, exceeding 50% on day three and reaching 100% on day 5 (Supplementary Figure 1A, Supplementary Table 1). Additionally, the surviving flies showed CCRT values higher than the control (Supplementary Figure 1B). The remaining temperatures resulted in minimal mortality, reaching an average of 10% at 8˚C and almost no mortality at 10˚C and 12˚C. These temperatures also lead to an improvement in the CCRT phenotype (Supplementary Figure 1B, Supplementary Table 2). Even though the effect of the remaining temperatures did not significantly differ from one another, the difference to control was largest when the acclimation temperature was 10˚C, and the duration was four days, which was chosen as the acclimation set-up for further experiments.

### Supplementary Methods

#### Cold Acclimation Preliminary Assays

To determine the temperature and duration of acclimation, we tested five time periods (1, 2, 3, 4, and 5 days) and four temperatures (6˚C, 8˚C, 10˚C, and 12˚C) using BKK12, BKK13, and KATH14, representing FastBKK, SlowBKK, and KATH populations, respectively. The ranges of temperature and duration were determined based on acclimation studies on other *Drosophila* species. The flies were controlled for age and the effect of acclimation was tested using the survival of acclimated flies and chill coma recovery time (CCRT) upon acclimation. Five replicates were used for each acclimation regime. On the day of emergence, 10 male flies were collected and placed into 50 ml vials with 5 ml food at room temperature. On the following day, test flies were flipped into new vials that were kept at the acclimation temperature and placed into cold incubators for 5-, 4-, 3-, 2-, and 1-day acclimation. The control flies were kept at rearing temperature (22 ± 1°C). After acclimation, the flies were placed back to room temperature for 24 hours for recovery, after which the dead flies were counted and removed from the vials, whereas the surviving flies were flipped into empty vials and placed on melting ice for 3 hours for CCRT measurement. At the end of 3 hours, the vials were placed into room temperature and the recovery time of each fly was measured. The temperature and duration that lead to minimal mortality and improved the CCRT were determined.

### Supplementary Figures

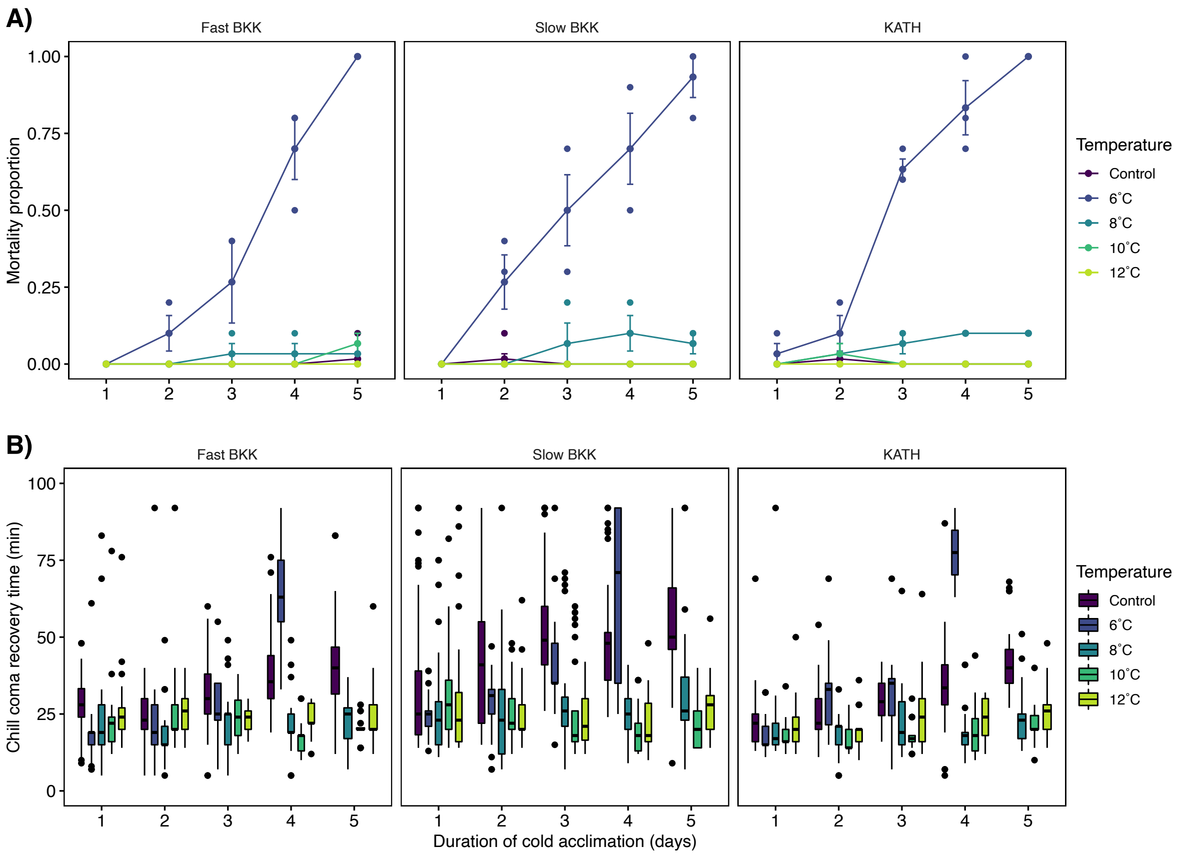


**Supplementary Figure 1:** Effect of duration of acclimation on A) mortality and B) chill coma recovery time (CCRT) phenotype. The lines that were used were representatives of the respective population and phenotype. Ten flies per replicate in five replicates were used for mortality measurements and dead flies were discarded. The remaining flies were used for chill coma recovery assays. Colored dots indicate proportional mortality of each replicate and black dots indicate outliers with high or low CCRT.

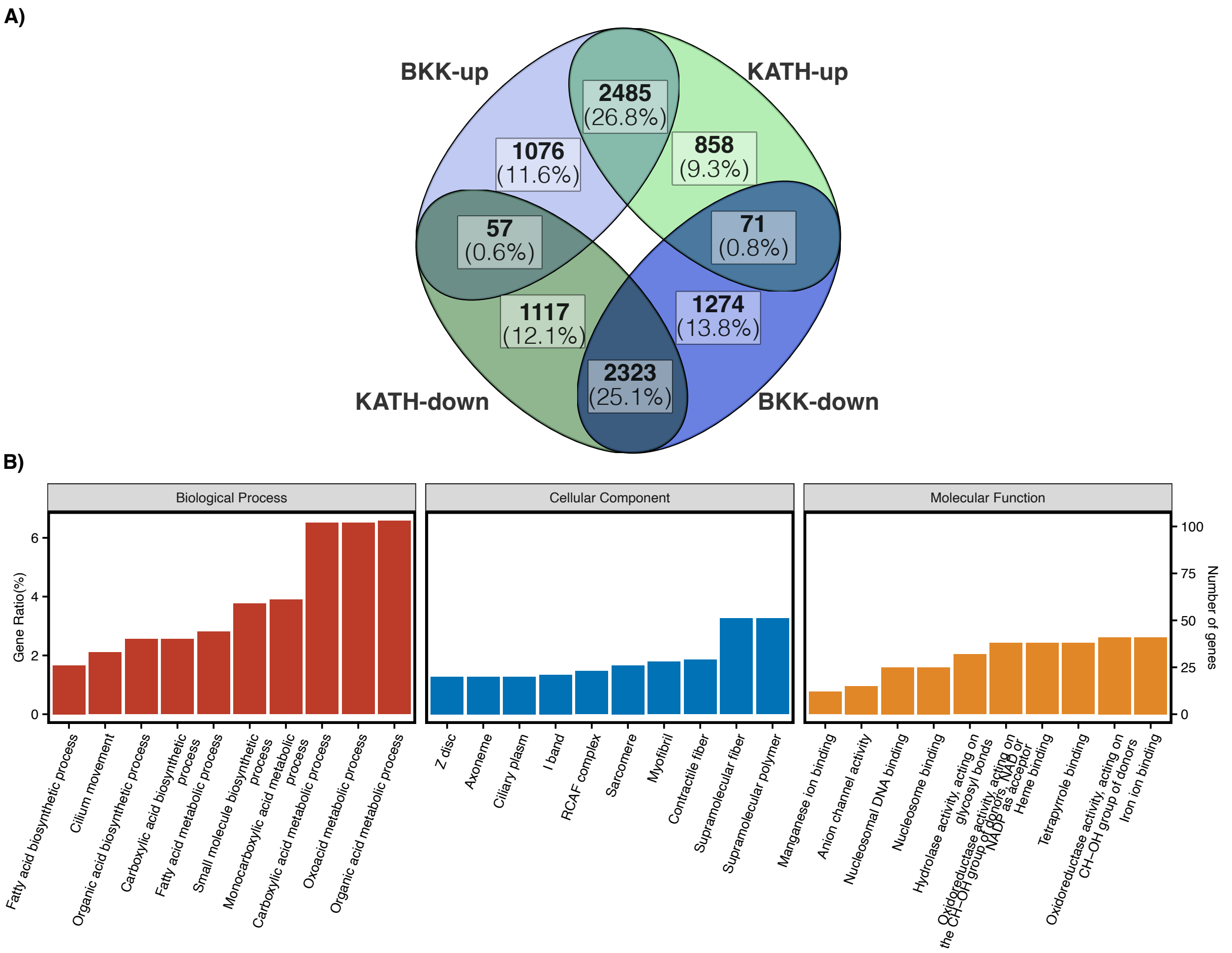


**Supplementary Figure 2:** A) Venn diagram with the number of differentially expressed (DE) genes specific to or shared between populations. Bangkok and Kathmandu populations exhibited both shared, as well as exclusive differentially expressed genes in response to acclimation. B) Gene ontology (GO) enrichment results for genes differentially expressed between the populations in response to cold acclimation treatment in the whole body. Biological processes (red bars), cellular components (blue bars), and molecular functions (yellow bars) are shown in separate panels.

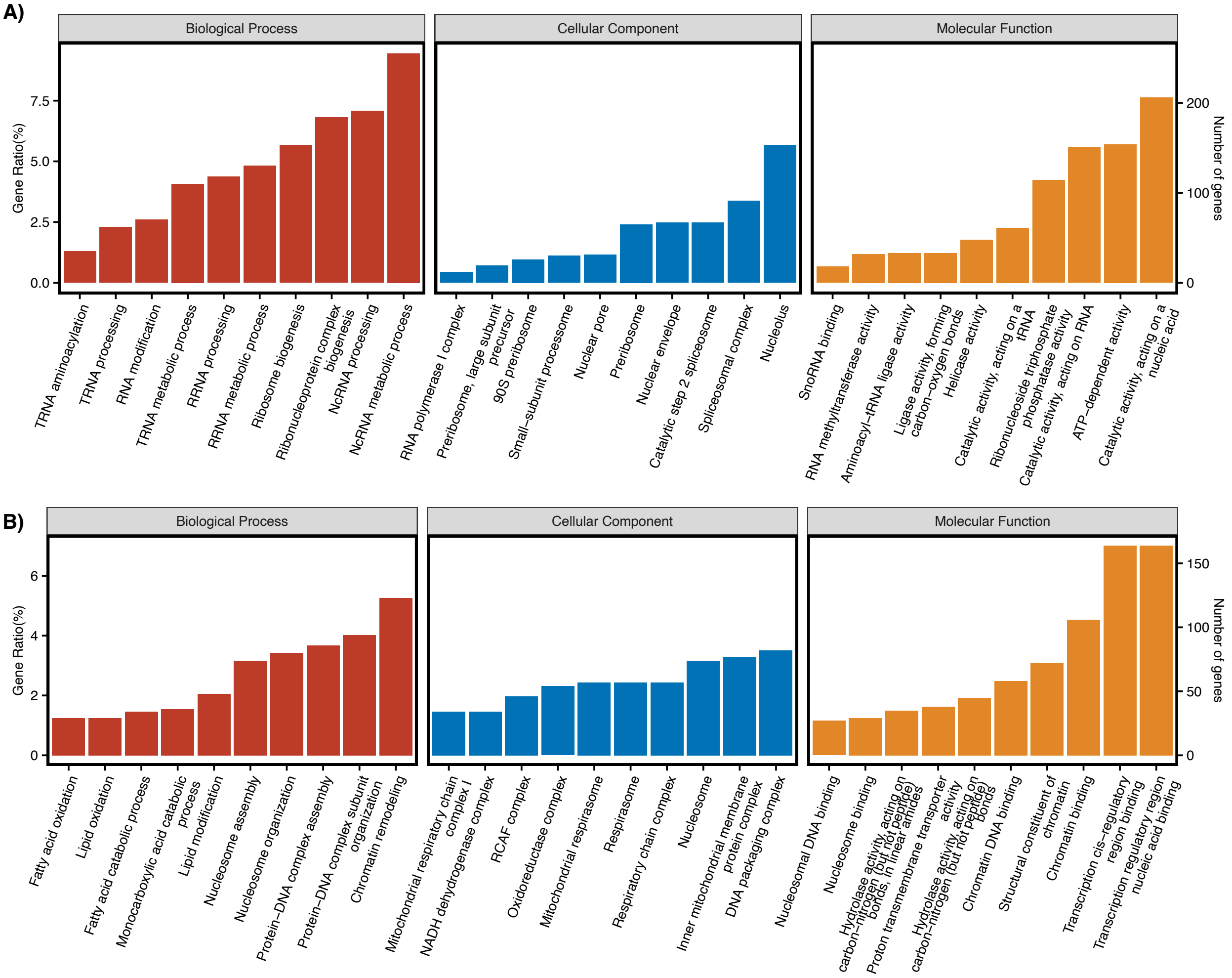


**Supplementary Figure 3:** Transcriptional response to cold acclimation in the Malpighian tubules. Gene enrichment analysis results for genes A) up-regulated and B) down-regulated in response to cold acclimation. Biological processes (red bars), cellular components (blue bars), and molecular functions (yellow bars) are shown in separate panels.

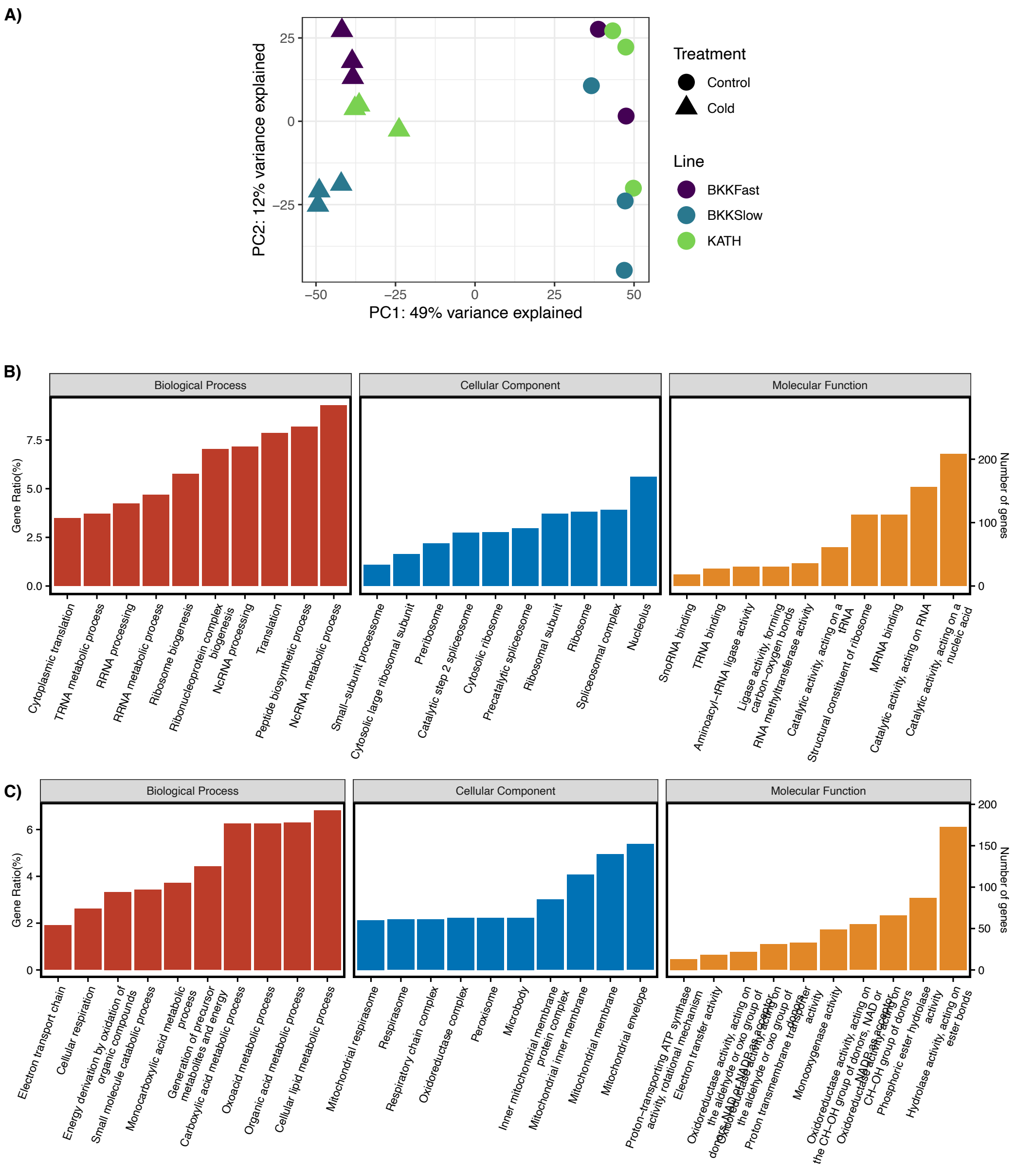


**Supplementary Figure 4:** Transcriptional response to cold acclimation in the hindguts. A) Principal component analysis (PCA) after the outlier was removed from the analysis. The samples are shown according to the pooled strains (color) and treatment (shape). FastBKK, SlowBKK, and KATH sample pools are represented by purple, blue, and green colors, respectively. The control samples are represented by circles and the cold acclimated samples are represented by triangles. Gene enrichment analysis results for genes B) up-regulated and C) down-regulated in response to cold acclimation. Biological processes (red bars), cellular components (blue bars), and molecular functions (yellow bars) are shown in separate panels. Cellular respiration and chitin metabolism play roles in the acclimation response of tolerant and sensitive strains in the hindgut.

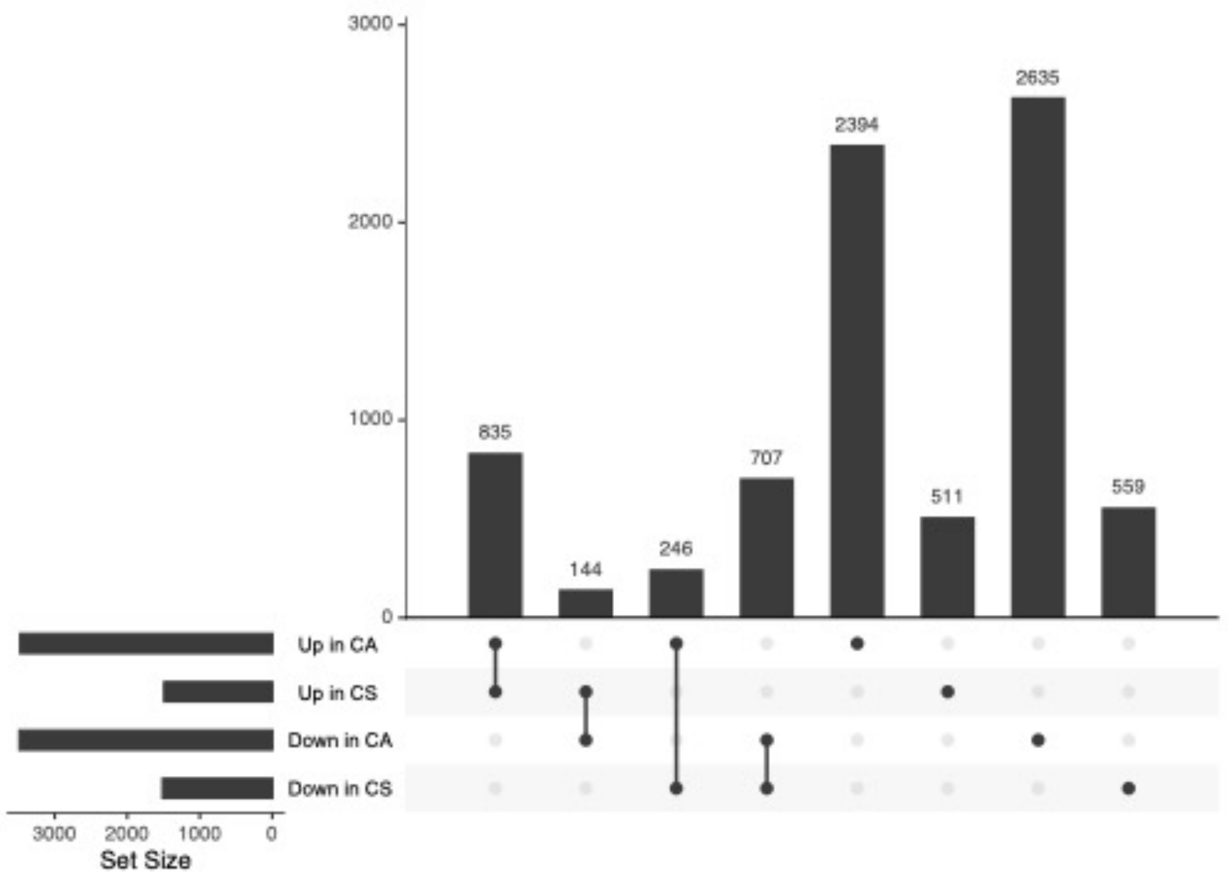


**Supplementary Figure 5:** Upset plot showing differentially expressed genes and their overlap between the cold shock (CS) and cold acclimation (CA) assays. Black, horizontal bars indicate the total number of differentially expressed genes that were significantly up- or down-regulated in a sample in response to cold acclimation. A black circle indicates that a sample and direction (up/down) combination is included in an intersection class. Intersection classes with a single circle are comprised of genes significantly differentially expressed in a single treatment and direction. Circles connected by a line indicate intersection classes comprised of multiple treatment and direction combinations. Vertical bars correspond to the number of genes in each intersection class.

### Supplementary Tables

**Supplementary Table 1:** Tukey's HSD test for proportional mortality after cold exposure.

| | 1-day | 2-day | 3-day | 4-day | 5-day |
|---|---|---|---|---|---|
| **6˚C** | e | d | c | b | a |
| **8˚C** | e | e | de | de | de |
| **10˚C** | e | e | e | e | e |
| **12˚C** | e | e | e | e | e |
| **control** | e | e | e | e | e |

**Note 1:** Samples that do not share any letter(s) are significantly different. *E.g.* Mortality upon 2-day exposure to 6˚C (**d**) significantly differs from mortality upon 3-day exposure to 6˚C (**c**), but not from mortality upon 3-day exposure to 8˚C (**de**).

**Supplementary Table 2:** Tukey's HSD test for  CCRT values upon acclimation.

| | 1-day | 2-day | 3-day | 4-day | 5-day |
|---|---|---|---|---|---|
| **8˚C** | def | ef | cdef | def | cde |
| **10˚C** | def | def | def | f | ef |
| **12˚C** | cde | def | def | def | cde |
| **control** | cd | c | b | b | a |

**Note 1:** Samples that do not share any letter(s) are significantly different. *E.g.* CCRT of 2-day control samples (**c**) significantly differs from CCRT of 3-day control samples (**b**), but not from CCRT of 1-day control samples (**cd**).
**Note 2:** Since the flies that survived the acclimation were used in the chill coma recovery assay, 6˚C acclimation was not included in the statistical test for CCRT due to high mortality (50-100%).

**Supplementary Table 3:** GO enrichment analysis results for differentially expressed genes exclusively in the whole body, Malpighian tubule, or hindgut samples in response to treatment.

**Supplementary Table 4:** GO enrichment analysis results for differentially expressed genes in the whole body, Malpighian tubule, and hindgut samples in response to treatment in opposite directions.

**Supplementary Table 5:** GO enrichment analysis results for differentially expressed genes in the whole body in response to treatment.

**Supplementary Table 6:** GO enrichment analysis results for differentially expressed genes in the whole body in response to treatment. The comparison of Kathmandu and Bangkok populations is shown. The column "Population" indicates which population showed positive log2FC value.

**Supplementary Table 7:** GO enrichment analysis results for differentially expressed genes in the whole body in response to treatment. The comparison of fast- and slow-recovering Bangkok lines is shown. The column "Line" indicates which line showed positive log2FC value.

**Supplementary Table 8:** GO enrichment analysis results for differentially expressed genes in the Malpighian tubules in response to treatment.

**Supplementary Table 9:** GO enrichment analysis results for differentially expressed genes in the Malpighian tubules in response to treatment. The comparison of Kathmandu and Bangkok populations is shown. The column "Population" indicates which population showed positive log2FC value.

**Supplementary Table 10:** GO enrichment analysis results for differentially expressed genes in the hindguts in response to treatment.

**Supplementary Table 11**: GO enrichment analysis results for differentially expressed genes in the hindguts in response to treatment. The comparison of fast and slow phenotypes is shown. The column "Phenotype" indicates which phenotype showed positive log2FC value.

**Supplementary Table 12:** GO enrichment analysis results for differentially expressed genes in the whole body in response to cold acclimation and cold treatment.

**Supplementary Table 13:** GO enrichment analysis results for differentially expressed genes in the whole body in response to cold acclimation or cold treatment.

**Supplementary Table 14:** GO enrichment analysis results for differentially expressed genes in the whole body in response to cold acclimation and cold treatment in opposite directions.
